# Dual projecting cells linking thalamic and cortical communication routes between the medial prefrontal cortex and hippocampus

**DOI:** 10.1101/2021.07.21.453279

**Authors:** Maximilian Schlecht, Maanasa Jayachandran, Gabriela E. Rasch, Timothy A. Allen

## Abstract

The interactions between the medial prefrontal cortex (mPFC) and the hippocampus (HC) are critical for memory and decision making and have been specifically implicated in several neurological disorders including schizophrenia, epilepsy, frontotemporal dementia, and Alzheimer’s disease. The ventral midline thalamus (vmThal), and lateral entorhinal cortex and perirhinal cortex (LEC/PER) constitute major communication pathways that facilitate mPFC-HC interactions in memory. Although vmThal and LEC/PER circuits have been delineated separately we sought to determine whether these two regions share cell-specific inputs that could influence both routes simultaneously. To do this we used a dual fluorescent retrograde tracing approach using cholera toxin subunit-B (CTB-488 and CTB-594) with injections targeting vmThal and the LEC/PER in rats. Retrograde cell body labeling was examined in key regions of interest within the mPFC-HC system including: (1) mPFC, specifically anterior cingulate cortex (ACC), dorsal and ventral prelimbic cortex (dPL, vPL), and infralimbic cortex (IL); (2) medial and lateral septum (MS, LS); (3) subiculum (Sub) along the dorsal-ventral and proximal-distal axes; and (4) LEC and medial entorhinal cortex (MEC). Results showed that dual vmThal-LEC/PER-projecting cell populations are found in MS, vSub, and the shallow layers II/III of LEC and MEC. We did not find any dual projecting cells in mPFC or in the cornu ammonis (CA) subfields of the HC. Thus, mPFC and HC activity is sent to vmThal and LEC/PER via non-overlapping projection cell populations. Importantly, the dual projecting cell populations in MS, vSub, and EC are in a unique position to simultaneously influence both cortical and thalamic mPFC-HC pathways critical to memory.

**Significance Statement:** The interactions between mPFC and HC are critical for learning and memory, and dysfunction within this circuit is implicated in various neurodegenerative and psychiatric diseases. mPFC-HC interactions are mediated through multiple communication pathways including a thalamic hub through the vmThal and a cortical hub through lateral entorhinal cortex and perirhinal cortex. Our data highlight newly identified dual projecting cell populations in the septum, Sub, and EC of the rat brain. These dual projecting cells may have the ability to modify the information flow within the mPFC-HC circuit through synchronous activity, and thus offer new cell-specific circuit targets for basic and translational studies in memory.

## 1. Introduction

Memory depends on bidirectional communication between the medial prefrontal cortex (mPFC) and hippocampus (HC) through (1) the ventral midline thalamus (vmThal) and (2) and lateral entorhinal cortex and perirhinal cortex (LEC/PER) (Dolleman-Van der Weel et al., 2019; Eichenbaum, 2017; Hoover & Vertes, 2007; Jin & Maren, 2015; Preston & Eichenbaum, 2013). At the cognitive and behavioral levels, the connections between mPFC and HC provide the interface for executive functions, working memory, and long-term memory integration (Eichenbaum, 2017; Eichenbaum et al., 1996; Hoover & Vertes, 2012; Jayachandran et al., 2019; Öngür & Price, 2000; Preston & Eichenbaum, 2013; Sigurdsson & Duvarci, 2016; Viena et al., 2018; Wirt & Hyman, 2017; W. Xu & Südhof, 2013). Rodent HC is considered homologous to other mammalian HC (Clark & Squire, 2013; Manns & Eichenbaum, 2006), while rodent mPFC is composed of agranular medial frontal cortex and shows some anatomical homologies to areas of the primate mPFC, but not granular lateral PFC (Allen & Fortin, 2013; Laubach et al., 2018; Passingham & Wise, 2012; Roberts & Clarke, 2019). Although the anatomy of the thalamic and cortical pathways in mPFC-HC interactions have been explored in detail, it is not known about whether the afferent regions shared by vmThal and LEC/PER arise from the same or different (non-overlapping) cell populations. Shared afferent cell populations have great potential to control the activity in the mPFC-HC circuit by synchronizing and/or gating the direction of activity flow through both thalamic and cortical communication pathways.

Generally, mPFC is thought to sit atop a cognitive system responsible for flexible behavior and executive functions (Berendse & Groenewegen, 1991; Eichenbaum, 1996; Guise & Shapiro, 2017; Laubach et al., 2018; Vertes, 2004; Vogt et al., 2013; Vogt & Paxinos, 2014; C. Xu et al., 2016). The HC, a core memory structure in the brain, has strong projections to mPFC with very few direct return projections from mPFC. In fact, return projections were only recently described by Malik et al., (2021) as a population of direct GABAergic projections from mPFC to inhibitory vasoactive intestinal polypeptide (VIP) interneurons in dorsal HC. Most mPFC communication with HC is polysynaptic. mPFC projections target vmThal, particularly lateral areas of the nucleus reuniens (RE, specifically periRE), and the parahippocampal region including LEC and PER. These cortical structures are notable for having bidirectional monosynaptic connections to one or more subfields of the HC proper (Bokor et al., 2002; Eichenbaum, 2017; Vertes, 2002; Wirt & Hyman, 2017) completing an HC to mPFC to HC loop.

RE, the largest region of vmThal, has a major role in cognitive tasks that place demands on spatial memory and/or executive function (Anderson et al., 2016; Cholvin et al., 2018; Van der Werf et al., 2002; Viena et al., 2018). The role of RE may be in modulating memory and setting up the constellations of cortical network modes best suited to the current behavioral and neural conditions. RE projections are a major source of thalamic afferents to HC targeting both excitatory and inhibitory neurons in the stratum lacunosum moleculare (slm) layer of CA1 (Cassel et al., 2013; Dolleman-Van der Weel & Witter, 2000; Mathiasen et al., 2019; McKenna & Vertes, 2004; Varela et al., 2014; Vertes, 2015; Vertes et al., 2007). Lateral RE (or periRE) receives extensive input from the entire dorsal-ventral extent of mPFC (layers V and VI), accompanied by afferents from Sub, cortex, and the basal forebrain (Griffin, 2015; Krout et al., 2002; McKenna & Vertes, 2004; Vertes et al., 2015). In addition to CA1, RE projections also innervate Sub (dorsal and ventral), mPFC, LEC, and PER (Vertes, 2006; Vertes et al., 2006; Wouterlood et al., 1990). These projections help define RE as a higher-order cortico-thalamo-cortical thalamic region at the diencephalic center of the mPFC-HC system (Dolleman-Van der Weel et al., 2019). Within HC, RE innervations of CA1 converge with EC layer III inputs in slm (temporoammonic pathway) synapsing on both excitatory and inhibitory neurons.

Long standing research has demonstrated a role for the parahippocampal region in learning, memory, and perception (Feinberg et al., 2012; Furtak et al., 2007; Hernandez et al., 2017; Kerr et al., 2007; Rodo et al., 2017). Within this region, LEC and PER serve as a site of massive cortical convergence providing one of the main communication hubs between mPFC and HC (Agster et al., 2016; Bartko et al., 2007b; Beckstead, 1979; Furtak et al., 2007; Hernandez et al., 2017; Hwang et al., 2018; Squire, 2009). This route is often described as part of the “what” pathway in memory circuitry (focused on here), as compared to MEC which is thought to comprise part of the “where” pathway (Knierim et al., 2014). LEC is a major input and output structure of the HC with convergent input from a variety of sensory and associational cortices (Canto et al., 2008; Knierim et al., 2006; Nilssen et al., 2019; Witter et al., 2017). Likewise PER, located along the rhinal sulcus (Burwell, 2001), is also involved in both memory (Suzuki et al., 1993) and higher-order sensory perception (Bartko et al., 2007a; Kent & Brown, 2012). Importantly, LEC and PER share mutual connections with mPFC (Agster & Burwell, 2009; Bedwell et al., 2015; Burwell & Amaral, 1998a; Sesack et al., 1989) and HC (Naber et al., 1999; Witter et al., 2000, 2006). PER, to a lesser degree than LEC, communicates directly with CA1 and is an important source of inputs and outputs for LEC itself. By contrast to thalamic mPFC-HC pathways through RE, LEC projects to all divisions of the hippocampal formation including dentate gyrus, CA1, CA3 as well as Sub (Amaral & Witter, 1989; Dolleman-Van der Weel & Witter, 1996; Witter, 2007; Witter et al., 1989).

Although it is known that the thalamic (mPFC-vmThal-HC) and cortical (mPFC-LEC/PER-HC) pathways make distinct contributions to memory (Barker et al., 2007; Barker & Warburton, 2008; Heimer-McGinn et al., 2017; Jayachandran et al., 2019; Kent et al., 2016; Kent & Brown, 2012; Paz et al., 2007; Ramanathan et al., 2018; Suzuki & Naya, 2014; Troyner et al., 2018; W. Xu & Südhof, 2013), the interactions between these two pathways is less studied, which have primarily focused on direct connectivity (Dolleman van der Weel, et al., 2019). Here we sought to identify outside cell populations that simultaneously communicate with vmThal and LEC/PER. To do this, we used a dual fluorescence-based retrograde tracing approach with cholera toxin subunit-B (CTB). We found that septum, Sub, and EC, but not mPFC or HC proper had dual vmThal-LEC/PER projecting cell populations.

## 2. Material and Methods

### 2.1. Animals

All animal experimental procedures were conducted in accordance with the Florida International University (FIU) Institutional Animal Care and Use Committee. Subjects were Long-Evans rats (n = 8; 4 females; Charles River Laboratories; weighing 250-350 g upon arrival). Tissue from four of the animals were previously used in Jayachandran et al., (2019) which reported mPFC (ACC, PL, and IL) pathways. Rats were individually housed and maintained on a 12 hr inverse light/dark cycle (lights off at 10 am). Rats had ad libitum access to food and water. Animals were named according to scheme NCLXXX (Neurocircuitry and Cognition Lab; XXX being an internal numbering scheme).

### 2.2. Micropipette pulling

Custom glass micropipettes were constructed with a laser-based micropipette puller (Sutter Instruments, P-2000) for Cholera toxin subunit-B (CTB) intracranial injections and designed to minimize the spread of the tracer. For LEC/PER injections, a taper length of about 6 – 8 mm was used. For vmThal injections, a taper length of 5 – 7 mm was used. The inner diameter for pipettes was between 80 – 100 µm and was accomplished using multiple heat cycles. The tip of the pipette was examined under a confocal microscope (Olympus FV1200) to verify accurate measurement and check for any visible damage. Prior to surgery, the tracer was loaded into each glass pipette and backfilled with mineral oil using a Hamilton syringe (Hamilton Company, Reno, NV).

### 2.3. Stereotaxic infusions of anatomical tracers

CTB tracers conjugated with two different Alexa Fluorophores (CTB-488 and CTB-594) were used as retrograde tracers. CTB was dissolved at a concentration of 1% in 0.06 M phosphate buffer. Animals were anesthetized with a combination of isoflurane (5% at beginning of the surgery, maintaining at 1-2%) and oxygen (800 mL/min). The head was fixed in a stereotaxic apparatus (David Kopf Instruments, Model 900), shaved clean, and betadine and isopropyl alcohol applied. Local injections of Marcaine were given prior to incision (7.5 mg/mL, 0.5 mL, s.c.) and the skull was exposed with a straight incision. Injection coordinates were calculated from bregma according to the Paxinos & Watson (2013) atlas. Bregma and lambda were leveled (±0.03 μm in the D/V plane) and burr holes were drilled overlying vmThal at a 10° angle to avoid the superior sagittal sinus; (A/P -1.8 mm, M/L -1.2 mm, D/V -6.85 mm) and LEC/PER (A/P - 6.0 mm, M/L -6.8 mm, D/V -6.5 mm) into the skull. Rats received 2.0 – 5.0 mL of Ringers solution throughout the surgery. Injections were performed using a Nanoject III microinjector with a glass micropipette (3.5” 3-000-203 G/X; Drummonds Scientific). Rats received unilateral injections of 0.3 µL into vmThal and 0.5 µL into LEC/PER at a flow rate of 1 nL/s. Some animals received CTB-488 in vmThal and CTB-594 in parahippocampal cortex, and some vice versa in case of any tracer differences. After complete tracer injections, the pipette was left untouched for 15 minutes to allow tracer diffusion. The pipette was removed carefully, and the incision site was sutured and dressed with Neosporin®. The rats were returned to a clean home cage and monitored until they woke. One day following surgery rats were administered a dose of Flunixin (50 mg/ml, 2.5 mg/kg, s.c.) and Neosporin® was reapplied. After a 2-week incubation period, rats were perfused.

### 2.4. Immunostaining

Rats were deeply anesthetized with isoflurane (5%) and transcardially perfused with 200 mL heparinized 0.1 M phosphate-buffered saline, followed by 200 ml of 4% paraformaldehyde (PFA, pH 7.4). Brains were post-fixed at least 24 – 48 hr in 4% PFA and then placed in a 30% sucrose solution for cryoprotection. Frozen brain sections were cut on a neuroanatomical research cryostat (Leica Biosystems, CM3050s; 40 μm, coronal plane) into three sets of immediately adjacent sections. To visualize tracer expression, as well as pipette tracts, slices were mounted on glass slides and cover slipped using Vectashield® antifade mounting medium (H-1200) with 4’,6-diamidino-2-phenylindole (DAPI). One set of tissue was used for fluorescence and confocal imaging, and the second set was used to conduct a Cresyl violet stain to use the local cytoarchitecture and identify the exact injection location. The third set of backup tissue was stored at -20 degrees in an ‘antifreeze’ soluton..

### 2.5. Region of interest (ROI) overlays

Overlays from the Paxinos and Watson stereotaxic brain atlas were used as a guide to outline the boundaries of the observable injection sites and regions of interest (ROIs). Immediately adjacent Cresyl violet sections were used for comparison and refinement based on cytoarchitectural features. Cresyl staining was performed on slide mounted tissue sections with 0.5% Cresyl violet acetate. The tissue was serially dehydrated with increasing concentrations of ethanol and coverslipped with Permount (Thermo Fisher Scientific) for brightfield imaging.

### 2.6. Confocal imaging

Confocal images were acquired with an Olympus Fluoview (FV1200) microscope with a 20x objective for cell quantification and a 60x and/or 100x objective for higher magnification images. Standard filter cubes for green fluorescence (excitation 470 nm; emission 525 nm), red fluorescence (excitation 545 nm; emission 605 nm), and DAPI (excitation 350 nm; emission 460 nm) were used. Depending on the ROI (mPFC, Sep, Sub, EC) an image size appropriate to each region image was captured. For each rat, a total of 68 ROI images were taken throughout the anteroposterior axis of each ROI, with a 200 µm distance between anterior and posterior parts. The number of images per ROI were as follows: mPFC, 40; Septum, 8; Sub, 8; and EC, 12. The ratio of CTB-488, CTB-594, and dual-labeled cells to the total number of DAPI cells was calculated for each ROI by subdivision. mPFC was quantified by layers (I-VI) at eight locations (ACC, dPL, vPL, and IL) from A/P ∼ 3.72 to ∼ 3.00 from Bregma. Sep was analyzed in medial (MS) and lateral divisions (LSd, LSi, LSv), as previously identified (Bokor et al., 2002; Canteras & Swanson, 1992), and double-labeling analysis was performed from A/P ∼ 0.60 to ∼ 0.36 from Bregma. Sub was analyzed in both dorsal and ventral portions, which was further subdivided into distal and proximal regions from A/P ∼ -6.12 mm to ∼ -6.48 mm from Bregma. Since the injection sites for LEC/PER could confound retrograde interpretations, ROIs for LEC and MEC were taken far enough away to account for possible tracer spread through extracellular diffusion. LEC and MEC was analyzed by layer (I-VI) from A/P ∼ -6.60 mm to ∼ -6.84 mm relative to Bregma. Quantification of single-target projecting cells and dual projecting cells were performed by two different experimenters and then averaged across.

### 2.7. Image processing and cell counts

Confocal images taken were saved in a proprietary Olympus file format (.oib). To speed up the processing of the images for quantification, a custom pipeline using CellProfiler (v4.0.7) was built. This pipeline helped merge three different imaging channels, by automatically combining all frames within an image and removed noise. Since DAPI is easily distinguishable compared to the retrograde tracers used, another pipeline was built to quantify the DAPI-stained nuclei automatically. We later compared several manual ROIs counts to the automatic CellProfiler counts to further validate the accuracy of our pipeline. The cell counts from Experimenter 1 and Experimenter 2, and their combined averages were significantly correlated with the results yielded by CellProfiler (CellProfiler vs. Experimenter 1, r = .982; CellProfiler vs. Experimenter 2, r = .983; CellProfiler vs. Average, r = .996; all p < .001).

### 2.8. Statistical analysis

All data are expressed as mean ± SEM, unless otherwise noted. Unpaired *t*-tests, ordinary one-way and two-way ANOVA were performed using GraphPad Prism 9.0 (GraphPad Software Inc.), and Microsoft Excel 2020 (Microsoft Corporation). All other image processing was applied equally across the field of view, and sampled using FIJI (ImageJ), and CellProfiler v4.0.7 with custom pipelines and (described above). Figures were constructed with Adobe Illustrator (version 25.3.1).

## 3. Results

### 3.1. Topographical distribution of injection sites

Eight animals received paired stereotaxic injections of CTB-488 and CTB-594 that were well localized to vmThal and the LEC/PER injections. Figure 1 shows the spread of the tracer throughout the anteroposterior axis of vmThal (Figure 1A-B) and LEC/PER (Figure 1C-D) for each animal. The center of the injection was marked as the brain section with greatest dorsoventral and mediolateral spread of the tracer (Figure 1Bii and 1Dii). This was often at the same location where the tip of the pipette showed slight tissue damage. Images were taken every 100-200 µm in the anterior (Figure 1Bi and 1Di) and posterior (Figure 1Biii and 1Diii) direction from the injection center, and matched with the closest atlas plate, to determine the extent of the tracer spread. The vmThal infusions centered in RE, with some diffusion was evident along the lateral wings of RE (or periRE), as well as an upward spread into other areas of vmThal including central medial thalamic nucleus (CM) and parts of the rhomboid thalamic nucleus (Rh). Three rats showed a small minority of spread into the anterior portion of the paraventricular thalamic nucleus (PVT), and the interanteromedial thalamic nuclei (IAM) (Figure 1A_i-iii_). The LEC/PER infusions were mostly localized to PER and dorsal LEC. Four rats showed some upward spread into temporal cortex following the path of damage from the glass injection pipette.

**Figure 1.**
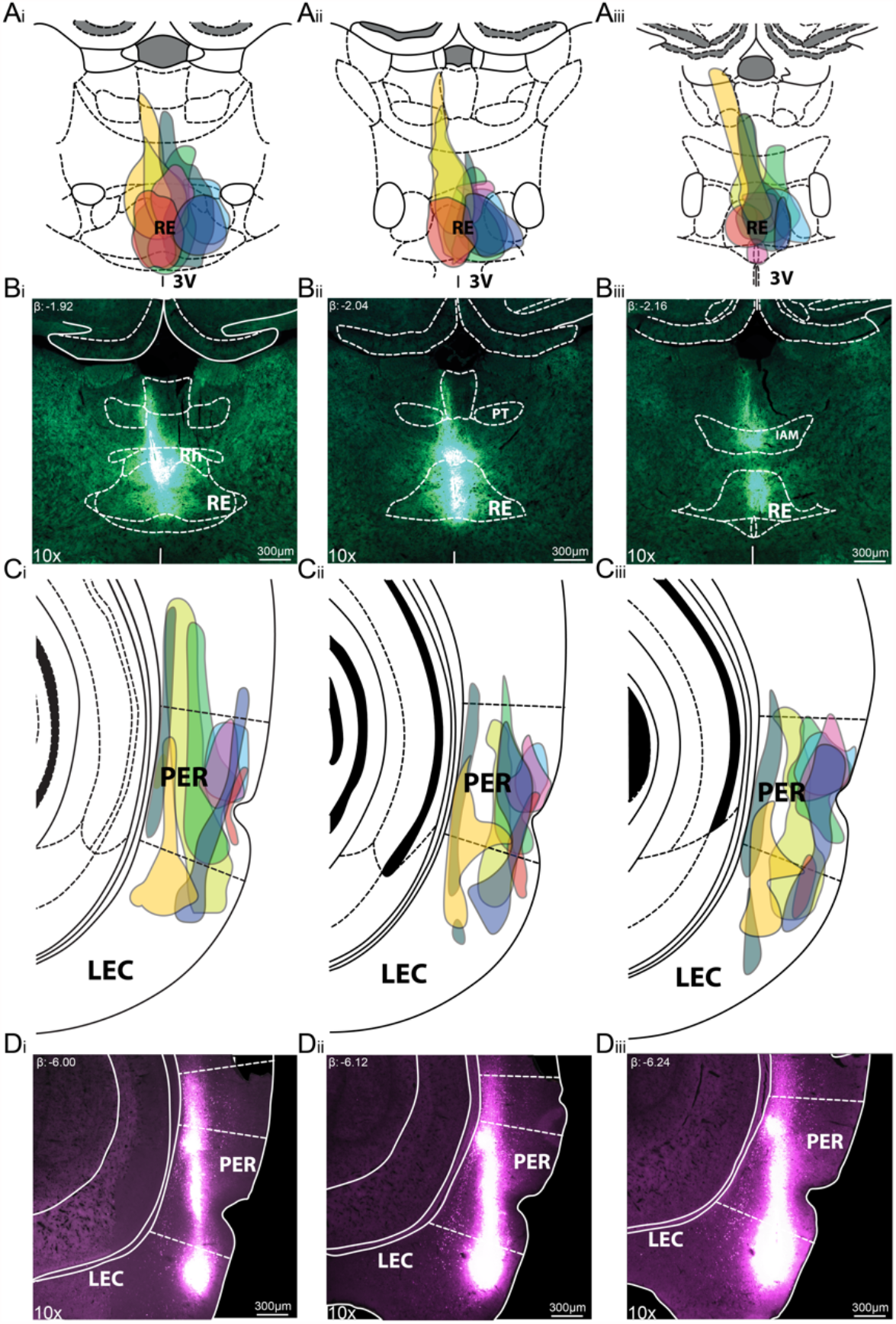
vmThal and LEC/PER CTB injection sites and spreads. **(A)** Mapping CTB in vmThal injection sites and spreads for all subjects in anterior (A_i_), medial (A_ii_), and posterior (A_iii_) sections, relative to the center of the injection. **(B)** Sample CTB vmThal injection site and spread in one subject showing anterior (B_i_), medial (B_ii_), and posterior (B_iii_) sections, relative to the center of the injection, light green subject shown. **(C)** Mapping CTB in LEC/PER injection sites and spreads for all subjects in anterior (C_i_), medial (C_ii_), and posterior (C_iii_) sections, relative to the center of the injection. **(D)** Sample CTB LEC/PER injection site and spread in one subject showing anterior (D_i_), medial (D_ii_), and posterior (D_iii_) sections, relative to the center of the injection, light green subject shown. The colors in A & C represent individual subjects.

In all eight rats, CTB-488 (green) and CTB-594 (magenta) cell body labeling was looked at in four ROIs: mPFC, Sep, Sub, and EC. mPFC was chosen for analysis here because this is known to control cognitive function through direct projections both vmThal and LEC/PER, although our initial examination did not reveal any individual cells projecting to both these areas (Jayachandran et al., 2019). Additional regions were chosen by searching the rest of the brain for regions containing individual cells labeled for both CTB-488 and CTB-594, which were found in Sep, Sub, and EC. Note, we also looked at CA1 in the HC, but did report that we did not observe dual labeled cell bodies in either the dorsal or ventral area.

### 3.2. mPFC

Rodent mPFC was analyzed in all major subdivisions (ACC, dPL, vPL, IL) across all layers (Layers I-VI) (Figure 2A). vmThal and LEC/PER-projecting cells were found in all mPFC subdivisions. vmThal cells were located primarily in layers V and VI of each subdivision, whereas LEC/PER-projecting cells were found primarily in layers II and III (Figure 2B). In mPFC, most of the vmThal-projecting cells are located in dPL, vPL, and IL with fewer in ACC. Generally, there were higher densities of vmThal-projecting neurons in mPFC. We did not find any dual vmThal-LEC/PER projecting cells in mPFC replicating earlier conclusions made in Jayachandran et al., 2019 (Figure 2D).

**Figure 2.**
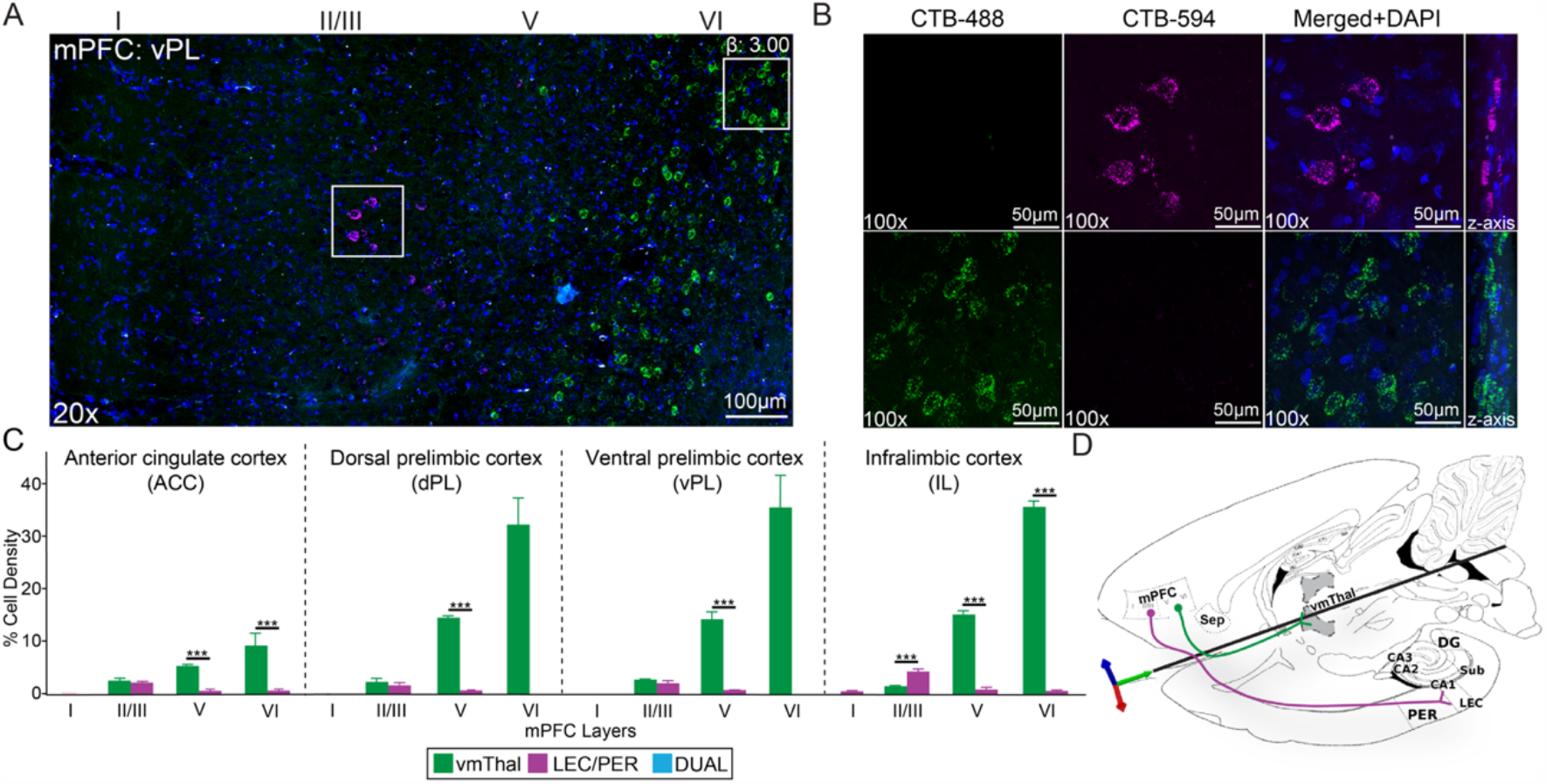
Medial prefrontal cortex: vmThal-projecting cells and LEC/PER-projecting cells. **(A)** 20x confocal montage from mPFC (specifically vPL) with vmThal-projecting cells (green, predominantly layers V-VI) and LEC/PER-projecting cells (magenta, predominantly layers II-V). No dual vmThal-LEC-PER projecting cells were found in mPFC. **(B)** 100x confocal images of white boxed areas in (A) showing CTB-488, CTB-594 and merged images with a z-stack on the right side. **(C)** Bar plot of the densities of ACC, dPL, vPL and IL cell projection populations analyzed across layers (I-VI). **(D)** Schematic representation of mPFC cell projection populations mapped onto combined sagittal-horizontal sections of the rat brain atlas. Data are represented as mean ± SEM. *p < 0.05; **p < 0.01; ***p < 0.001.

In ACC, there was a significant difference in the densities of vmThal-projecting neurons across layers I (0.587 ± 0.055%), II/III (1.794 ± 0.086%), V (5.204 ± 0.109%) and VI (9.129 ± 2.331%) tested using a one-way ANOVA F_(3,28)_ = 11.760, p < 0.0001. The highest density of vmThal-projecting cells was consistently found in layers V and VI. Very few vmThal-projecting neurons were found in shallow layers of ACC. There also was a significant difference in LEC/PER-projecting cells across layers I (0%), II/III (1.443 ± 0.185%), V (0.449 ± 0.339%) and VI (0.500 ± 0.313%) tested using a one-way ANOVA F_(3,28)_ = 5.952, p = 0.0028 (Figure 2C). Most LEC/PER-projecting cells were confined to layers II/III (Figure 2A) in ACC, in contrast to vmThal-projecting cells, thus demonstrating there is a significant interaction effect between the projection target and cortical layer (F_(3,56) projection x layer_ = 11.60, p < 0.0001, interaction effect; F_(1,56) projection_ = 33.340, p < 0.0001, main projection effect; F_(3,56) layer_ = 11.420, p < 0.0001, main layer effect).

In dPL, there was also significant difference in cell densities of vmThal-projecting neurons across layers I (0%), II/III (2.253 ± 0.615%), V (14.086 ± 0.379%), and VI (31.351 ± 4.929%) tested using a one-way ANOVA F_(3,28)_ = 33.180, p < 0.0001 (Figure 2C). The highest density of vmThal-projecting cells was consistently found in layers V and VI. Few vmThal-projecting neurons were observed in shallow layers of dPL, with none being found in layer I. There was also a significant difference in LEC/PER-projecting cells across layers I (0%), II/III (1.585 ± 0.507%), V (0.626 ± 0.106%) and VI (0%) tested using a one-way ANOVA F_(3,28)_ = 8.332, p < 0.0001 (Figure 2C). Most of the LEC/PER-projecting cells were restricted to layers II/III (Figure 2A) in dPL, in comparison to vmThal-projecting cells (F_(3,56) projection x layer_ = 34.40, p < 0.0001, interaction effect; F_(1,56) projection_ = 82.420, p < 0.0001, main projection effect; F_(3,56) layer_ = 31.430, p < 0.0001, main layer effect).

vPL showed a significant difference in densities of vmThal-projecting neurons across layers I (0%), II/III (2.838 ± 0.115%), V (14.237 ± 1.386%), and VI (35.366 ± 6.055%) tested using a one-way ANOVA F_(3,28)_ = 26.730, p < 0.0001. The highest density of vmThal-projecting cells was also consistently found in layers V and VI. Few vmThal-projecting neurons were found in shallow layers of vPL. A significant difference in LEC/PER-projecting cells was observed across layers I (0%), II/III (2.173 ± 0.461%), V (0.843 ± 0.059%), and VI (0.055 ± 0.053%) using a one-way ANOVA F_(3,28)_ = 18.750, p < 0.0001 (Figure 2C). Most LEC/PER-projecting cells were confined to layers II/III (Figure 2A) in vPL, in contrast to vmThal-projecting cells (F_(3,56) projection x layer_ = 28.070, p < 0.0001, interaction effect; F_(1,56) projection_ = 62.790, p < 0.0001, main projection effect; F_(3,56) layer_ = 25.300, p < 0.0001, main layer effect).

In IL, there was also a significant difference in the densities of vmThal-projecting neurons across layers I (0%), II/III (1.088 ± 0.305%), V (14.555 ± 0.657%) and VI (34.269 ± 1.828%) tested using a one-way ANOVA F_(3,28)_ = 263.400, p < 0.0001 (Figure 2C). The highest density of vmThal-projecting cells was repeatedly found in layers V and VI. Very few vmThal-projecting neurons were found in shallow layers of IL. There also was a significant difference in LEC/PER-projecting cells across layers I (0.168 ± 0.162%), II/III (4.108 ± 0.363%), V (0.785 ± 0.277%), and VI (0.118 ± 0.110%) tested using a one-way ANOVA F_(3,28)_ = 58.410, p < 0.0001. Most LEC/PER-projecting cells were confined to layers II/III (Figure 2A) in IL, in contrast to vmThal-projecting cells (F_(3,56) projection x layer_ = 280.100, p < 0.0001, interaction effect; F_(1,56) projection_ = 486.000, p < 0.0001, main projection effect; F_(3,56) layer_ = 222.200 , p < 0.0001, main layer effect).

### 3.3. Septum

Septum was analyzed in all major atlas-defined subdivisions including the lateral septum dorsal (LSd), lateral septum ventral (LSv), lateral septum intermediate (LSi) (Figure 3A) and medial septum (MS) (Figure 3B). vmThal and LEC/PER-projecting cells were found in all septal subdivisions, with the vast majority being vmThal-projecting cells. Importantly, all subdivisions showed the presence of dual vmThal-LEC/PER projecting cells, but these were predominantly located in MS (Figure 3D).

**Figure 3.**
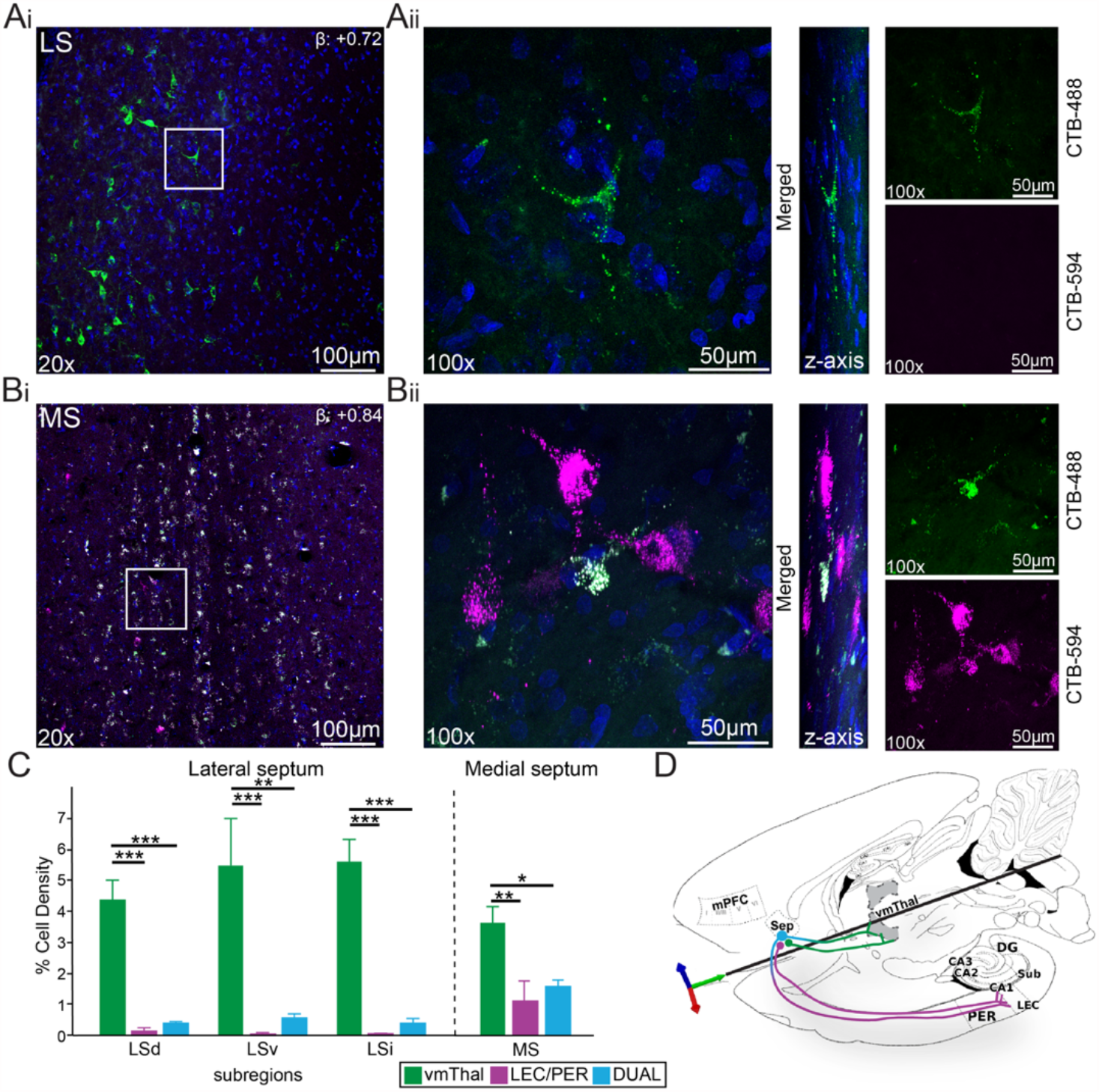
Septum: vmThal-projecting cells, LEC/PER-projecting cells, and dual vmThal-LEC/PER projecting cells. **(A**_**i**_) 20x confocal montage from LS showing vmThal-projecting cells. LEC/PER-projecting cells and dual vmThal-LEC/PER projecting cells were not frequently observed. **(A**_**ii**_) 100x confocal image (merged channels) of the whited boxed area in (A_i_) with a z-stack in the middle, and CTB-488 and CTB-594 channels separated on the right side. **(B**_**i**_) 20x confocal montage from MS showing abundant dual vmThal-LEC/PER projecting cells (cyan), including scattered vmThal-projecting cells (green) and LEC/PER-projecting cells (magenta). **(B**_**ii**_) 100x confocal image (merged channels) of the whited boxed area in (B_i_) with a z-stack in the middle, and CTB-488 and CTB-594 channels separated on the right side. **(C)** Bar plot of the densities of LSd, LSv, LSi, and MS cell projection populations. **(D)** Schematic representation of MS cell projection populations mapped onto combined sagittal-horizontal sections of the rat brain atlas. Data are represented as mean ± SEM. *p < 0.05; **p < 0.01; ***p < 0.001

In LS, there was a consistent population of vmThal-projecting cells across all subdivisions (LSd: 4.384 ± 0.627%; LSv: 5.497 ± 1.502%; LSi: 5.602 ± 0.721%). There was no significant difference in the prevalence of vmThal-projecting cells across the lateral subdivisions, including MS (one-way ANOVA: F_(3,28)_ = 1.027, p = 0.396 (Figure 3C). Very few LEC/PER projections were identified in lateral regions of septum (overall LS: 0.040%-0.25%; LSd: 0.143 ± 0.103%; LSv: 0.036 ± 0.035%; LSi: 0.061± 0.005%). vmThal-projecting cells were significantly denser compared to LEC/PER-projecting cells within each LS subregions (LSd, F_(2,21)_ = 42.190, p < 0.0001; LSv, F_(2,21)_ = 11.970, p = 0.0003; LSi, F_(2,21)_ = 53.110, p < 0.0001; Figure 3C).

In MS, there were fewer vmThal-projecting cells (3.646 ± 0.497%) compared to all LS subdivisions (Figure 3C). On the other hand, lateral septum did not have a lot of LEC/PER-projecting cells, whereas MS has a considerable amount (Figure 3C). Overall, the highest % of LEC/PER-projecting cells were found in MS (MS overall: 1.130 ± 0.642%). However, no significant differences were identified between all septum subdivisions when comparing LEC/PER-projecting cell populations (one-way ANOVA: F_(3,28)_ = 2.618, p = 0.0706). Within MS, vmThal-projecting cells continued to be significantly denser compared to LEC/PER-projecting cells (one-way ANOVA: F_(2,21)_ = 7.736, p = 0.0032).

A population of cells that simultaneously project to both vmThal and LEC/PER were discovered in MS (1.592 ± 0.211%; Figure 3D). Dual vmThal-LEC/PER-projecting cells in LS were found but accounted for less than one percent of total cells. In order to test whether the septal subdivision and projection population were closely related, an ordinary two-way ANOVA was conducted testing whether there is an interaction effect. While there was no significant interaction effect, a significant projection effect is observed (F_(6,84) projection x subdivision_ = 2.070, p = 0.0654; F_(2,84) projection_ = 73.850, p < 0.0001; F_(6,84) subdvision_ = 0.424, p = 0.736).

### 3.4. Subiculum

Subiculum was analyzed in all major subdivisions throughout, including dorsal (dSub; Figure 4Ai-ii) and ventral subiculum (vSub; Figure 4Bi-ii). Given the distinct structure of subiculum, cells were analyzed along the proximodistal axis, thus the major atlas subdivision of distal and proximal were applied onto both dSub and vSub. vmThal and LEC/PER-projecting cells were found in all subiculum subdivisions, with the majority being vmThal-projecting cells. Interestingly, LEC/PER-projecting cells were only found in vSub. Nonetheless, all subdivisions showed the presence of dual vmThal-LEC/PER-projecting cells, but these were also predominantly located in vSub.

**Figure 4.**
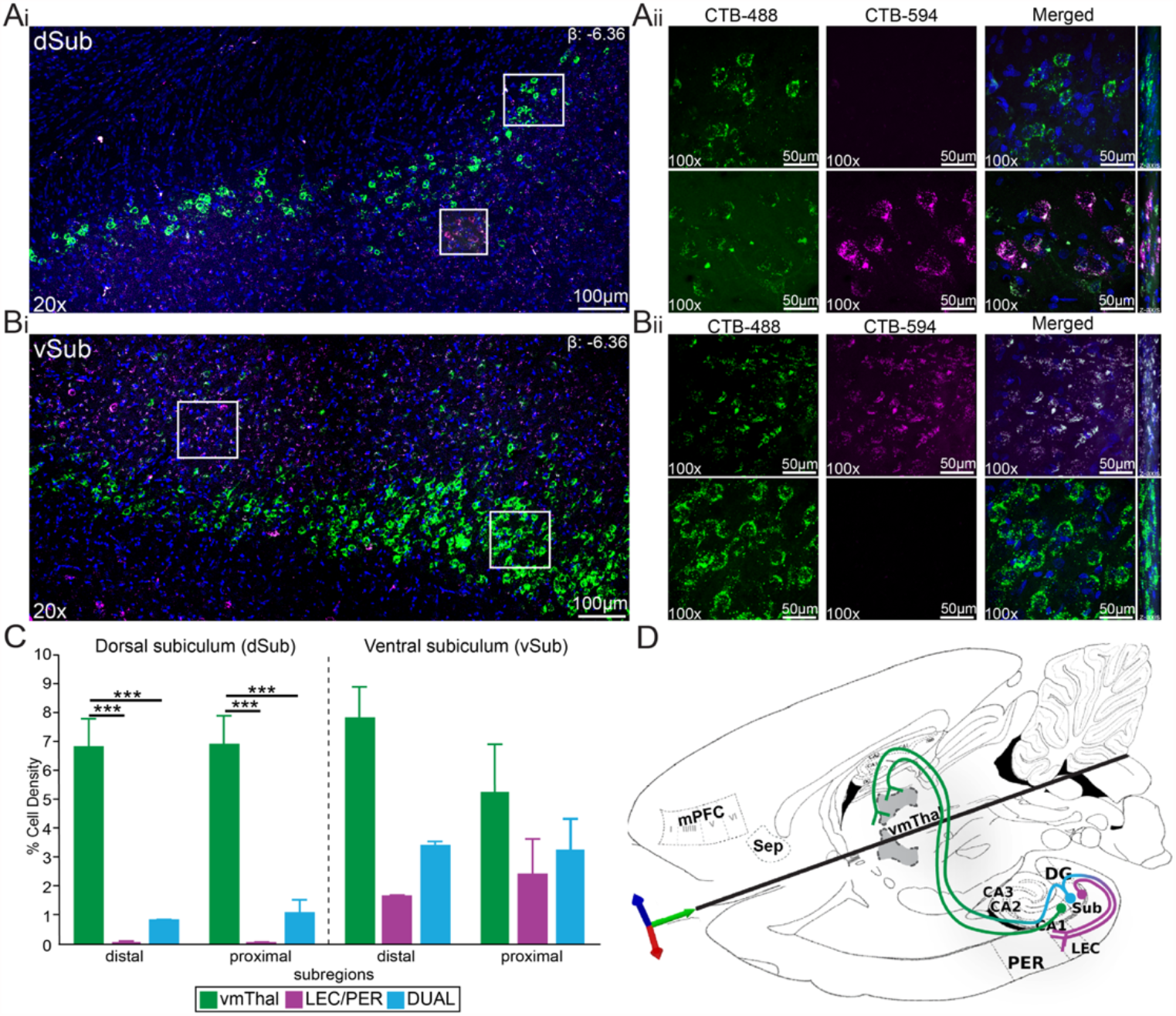
Subiculum: vmThal-projecting cells, LEC/PER-projecting cells, and dual vmThal-LEC/PER projecting cells. **(A**_**i**_) 20x confocal montage from dSub showing vmThal-projecting cells (green, predominantly pyramidal cell layer) and dual vmThal-LEC/PER projecting cells (cyan, polymorphic layer). LEC/PER projecting cells (magenta) were not commonly observed. **(A**_**ii**_) 100x confocal image of the whited boxed area in (A_i_) separated by channels for CTB-488 (green) and CTB-594 (magenta), and merged with a z-stack the right side. **(B**_**i**_) 20x confocal montage from dSub showing vmThal-projecting cells (green, predominantly pyramidal cell layer), LEC/PER projecting cells (magenta, all layers), and dual vmThal-LEC/PER projecting cells (cyan, polymorphic layer). **(B**_**ii**_) 100x confocal image of the whited boxed area in (B_i_) separated by channels for CTB-488 (green) and CTB-594 (magenta), and merged with a z-stack the right side. **(C)** Bar plot of the densities of dSub and vSub cell projection populations. **(D)** Schematic representation of MS cell projection populations mapped onto combined sagittal-horizontal sections of the rat brain atlas. Data are represented as mean ± SEM. *p < 0.05; **p < 0.01; ***p < 0.001.

In dSub, there was a continuous population of vmThal-projecting cells along the proximodistal axis (distal dSub: 6.886 ± 0.912%; proximal dSub: 6.943 ± 0.956%). There was no significant difference in the prevalence of vmThal-projecting cells across the dorsal subdivisions (unpaired t-test: t_(14)_ = 0.043, p = 0.962; Figure 4C). Very few LEC/PER-projecting cells were identified in the dorsal subdivisions of subiculum (distal dSub: 0.032 ± 0.018%; proximal dSub: 0.008 ± 0.009%). Importantly, a dual vmThal-LEC/PER-projecting cell population was found predominantly in proximal dSub (1.151 ± 0.390%), while in distal dSub (0.829 ± 0.031%), the population made up less than 1% (unpaired t-test: t_(14)_ = 0.823, p = 0.424; Figure 4C). In order to test whether the dSub subdivision and projection populations were closely related a two-way ANOVA was conducted testing for an interaction effect, but no significant interaction effect was observed (F_(2,42) projection x subdivision_ = 0.052, p = 0.949). However, a projection-specific effect was observed, in which vmThal-projecting cells were significantly denser than LEC/PER-projecting cells and dual vmThal-LEC/PER-projecting cells and evident in every subdivision (F_(2,42) projection_ = 88.000, p < 0.0001; F_(1,42) subdivision_ = 0.067, p = 0.798).

In vSub, vmThal-projecting cells were identified along the proximodistal axis (distal vSub: 7.845 ± 1.050%; proximal vSub: 5.288 ± 1.628%). There was no significant difference in the prevalence of vmThal-projecting cells across the ventral subdivisions (unpaired t-test: t_(14)_ = 1.320, p = 0.208; Figure 4C). A larger population of LEC/PER-projecting cells were identified in the ventral subdivisions of subiculum (distal vSub: 1.681 ± 0.019%; proximal vSub: 2.408 ± 1.236%). We then tested the differences of LEC/PER-projecting cells between dSub and vSub subdivisions and found subiculum projects almost exclusively to the parahippocampal cortices via a ventral pathway (one-way ANOVA: F_(3,28)_ = 3.807, p = 0.021). A much larger dual vmThal-LEC/PER-projecting cell population was identified in ventral subdivisions of both distal vSub (3.436 ± 0.104%) and proximal vSub (3.247 ± 1.091%; unpaired t-test: t_(14)_ = 0.173, p = 0.865). In a like manner, dual vmThal-LEC/PER-projecting cells from subiculum also project more through ventral pathways (one-way ANOVA: F _(3,28)_ = 8.524, p < 0.0001; Figure 4D).

### 3.5. Entorhinal Cortex

EC was subdivided in major atlas defined subdivisions into lateral (LEC) and medial (MEC) regions for cell quantification (distant from CTB injection sites). Each region was further subdivided according to layers (I-VI) found in the literature (Burwell & Amaral, 1998b; Insausti et al., 1997; Save & Sargolini, 2017; Sewards & Sewards, 2003; Witter et al., 2017). vmThal and LEC/PER-projecting cells were found in all EC subdivisions. vmThal-projecting cells were located primarily in shallow layers II/III in both LEC and MEC (Figure 5A), whereas LEC/PER-projecting cells were found primarily in layers II/III, V, and VI. Generally, there were higher densities of LEC/PER-projecting cells in both LEC and MEC. Interestingly, dual vmThal-LEC/PER-projecting cells were found almost exclusively in shallow layers II/III of each subdivision (Figure 5B and 5D).

**Figure 5.**
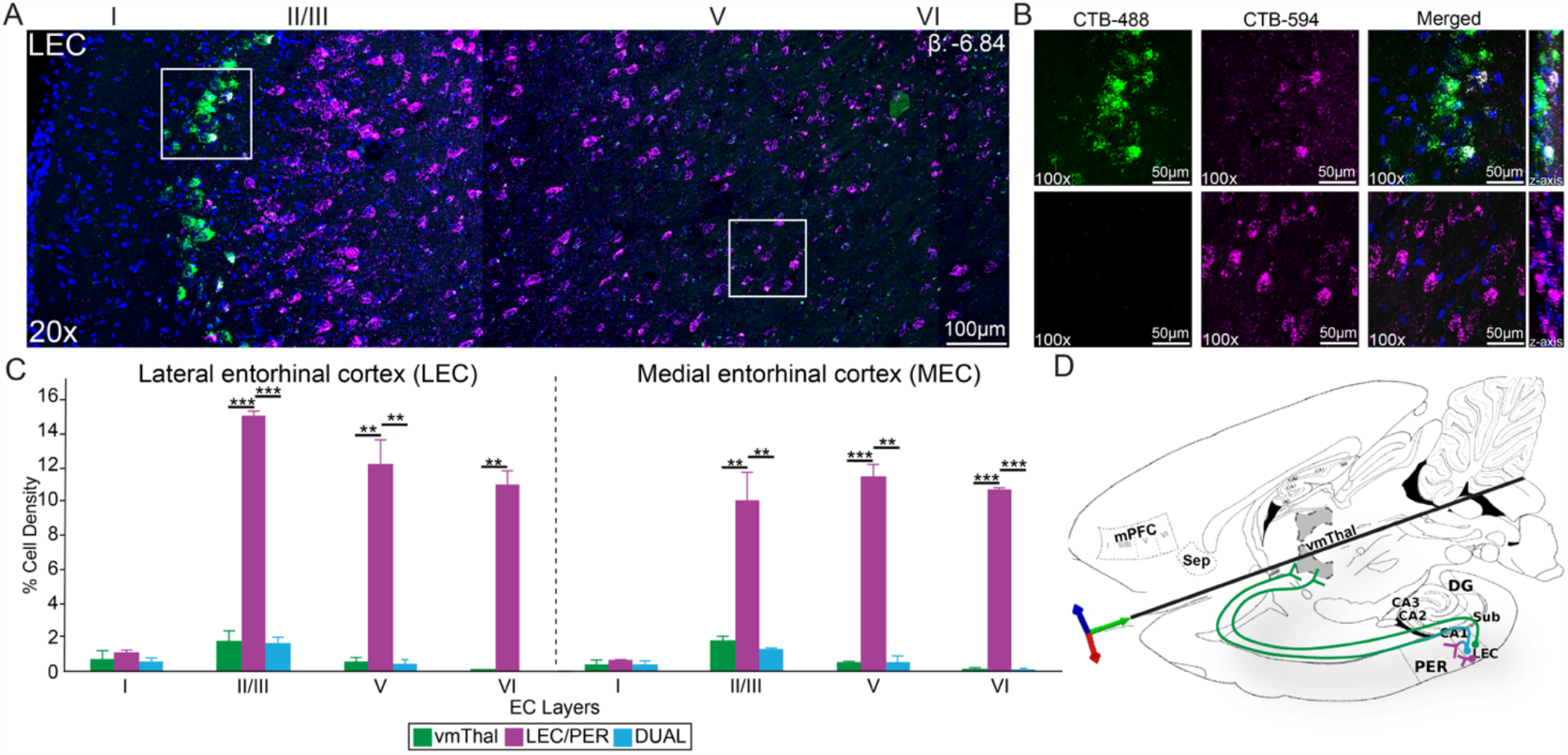
Entorhinal Cortex: vmThal-projecting cells, LEC/PER-projecting cells, and dual vmThal-LEC/PER projecting cells. **(A)** 20x confocal montage of LEC with vmThal-projecting cells (green; layer II), while LEC/PER-projecting cells (magenta; layers II-V), and dual vmThal-LEC/PER projecting cells (cyan; deep layer II). MEC not shown, but had a similar pattern. **(B)** 100x confocal images from the white boxed areas in (A) showing CTB-488, CTB-594, and merged images with a z-stack on the right. **(C)** Bar plot of the densities of LEC and MEC cell projection populations analyzed across layers (I-VI). **(D)** Schematic representation of LEC cell projection populations. Data are represented as mean ± SEM. *p < 0.05; **p < 0.01; ***p < 0.001.

In LEC, no significant differences were identified between vmThal-projecting cells across layers I (0.699 ± 0.513%), II/III (1.749 ± 0.628%), V (0.509 ± 0.297%), and VI (0.088 ± 0.003%) tested using a one-way ANOVA F_(3,28)_ = 2.669, p = 0.067. The highest density of vmThal-projecting cells was found in the shallowest layers of II/III, while very few vmThal-projecting cells were found in deeper layers of LEC (Figure 5A and 5C). There also was a significant difference in LEC/PER-projecting cells across layers I (1.135 ± 0.113%), layers II/III (15.091 ± 0.234%), layer V (12.170 ± 1.464%), and layer VI (10.955 ± 0.854%) tested using a one-way ANOVA F_(3,28)_ = 49.840, p < 0.0001.

A population of cells that simultaneously project to both vmThal and LEC/PER were discovered in LEC mainly in layers II/III (1.606 ± 0.405%), but also a small percentage in layer I (0.595 ± 0.188%), and layer V (0.405 ± 0.271%), with none in layer VI (one-way ANOVA: F_(3,28)_ = 6.825, p = 0.001; Figure 5D). Most LEC/PER-projecting cells were found in layers II/III, V, and VI in contrast to vmThal-projecting cells and dual vmThal-LEC/PER-projecting cells, thus supporting a interaction effect between projecting population and cortical layer (F_(6,84) projection x layer_ = 34.600, p < 0.0001, interaction effect; F_(2,84) projection_ = 336.600, p < 0.0001, main projection effect; F_(3,84) layer_ = 44.730, p < 0.0001, main layer effect). MEC showed significant differences in the densities of vmThal-projecting cells across layers I (0.356 ± 0.311%), II/III (1.845 ± 0.232%), V (0.552 ± 0.044%), and VI (0.189 ± 0.039%) tested using a one-way ANOVA F_(3,28)_ = 14.730, p < 0.0001. The highest density of vmThal-projecting cells continued to be found in the shallowest areas of layers II/III. There also was a significant difference in densities of LEC/PER-projecting cells across layers I (0.676 ± 0.016%), II/III (10.142 ± 1.593%), V (11.534 ± 0.672%), and VI (10.769 ± 0.052%) tested using a one-way ANOVA F_(3,28)_ = 34.780, p < 0.0001. Lastly, significant differences in densities of dual vmThal-LEC/PER projecting cells persisted across layers I (0.464 ± 0.155%), II/III (1.364 ± 0.017%), V (0.603 ± 0.318%), and VI (0.08%), tested using a one-way ANOVA F_(3,28)_ = 4.197, p = 0.014. An interaction effect was observed between the projection targets and cortical layer in MEC (F_(6,84) projection x layer_ = 29.340, p < 0.0001, interaction effect; F_(2,84) projection_ = 270.000, p < 0.0001, main projection effect; F_(3,84) layer_ = 35.620, p < 0.0001, main layer effect).

## 4. Discussion

### 4.1. Summary of main findings

In this study, we looked for cell populations that project to both the vmThal (targeting RE) and the LEC/PER using a dual fluorescence retrograde tracing approach (CTB-488 and CTB-594). This approach unambiguously identifies dual projecting cells through the colocalization of both CTB tracers (488+ and 594+ cells) in projection cells using optical sections analysis via confocal microscopy. This approach identified three key populations of dual vmThal-LEC/PER-projecting cells in (1) MS, (2) Sub, and (3) EC, summarized in Figure 6. First, we found that dual vmThal-LEC/PER-projecting cells were prominent in MS. The dual projecting MS cells were interspersed among vmThal-projecting and LEC/PER-projecting cells, with the former more abundant. While there were dual projecting cells in LS, they were a minority population by comparison to those in MS. LEC/PER-projecting cells were also found in LS but this was the least abundant cell projection type. Second, dual vmThal-LEC/PER-projecting cells were identified throughout the subiculum, with the highest levels observed in vSub (proximal and distal). Similar to MS and LS, vmThal-projecting cells were more abundant in dSub, while LEC/PER-projecting cells were a minority population in dSub. LEC/PER-projecting cells were prominent in vSub. Third, both LEC and MEC contained dual vmThal-LEC/PER-projecting cells, which were almost exclusively restricted to layer II/III (more specifically Layer IIb). vmThal-projecting cells were found in layers II/III, and V of both LEC and MEC, but most prominent in layer II/III. LEC/PER-projecting cells were found in layers II/III, V, and VI, but always underneath the vmThal-projecting layer. This pattern was similar in both MEC and LEC. While the cell populations targeting vmThal or LEC/PER have been described separately in great detail elsewhere (Agster & Burwell, 2009; Blackstad, 1956; Burwell & Amaral, 1998a; Canto et al., 2008; Griffin, 2015; McKenna & Vertes, 2004; Vertes et al., 2006; Wouterlood et al., 1990), the dual vmThal-LEC/PER-projecting cell populations have not been reported.

**Figure 6.**
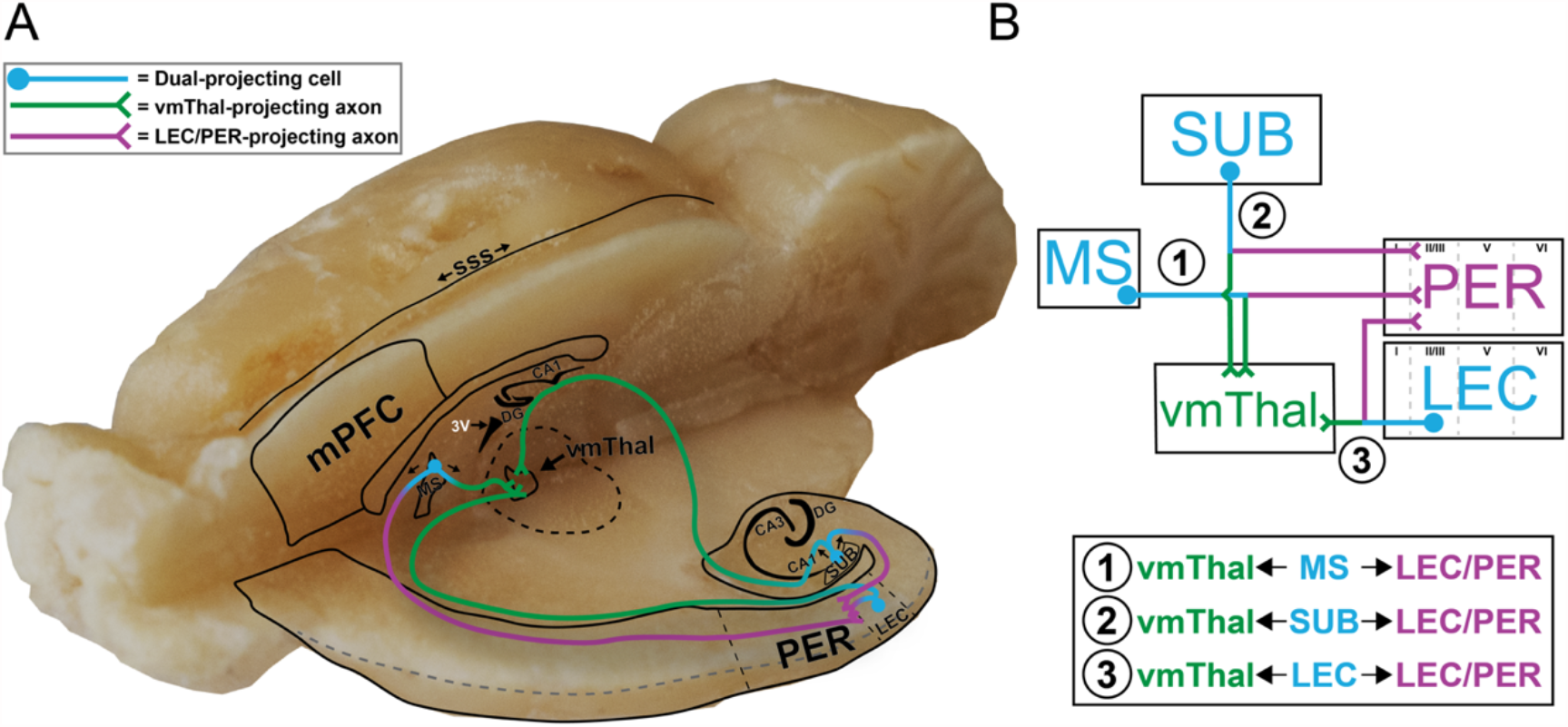
Dual vmThal-LEC/PER projecting cell populations in the mPFC-HC system. **(A)** A picture of a rat brain with a saggital-horizontal wedge removed and a model of three distinct dual vmThal-LEC/PER cell populations superimposed: (1) MS, (2) Sub, and (3) EC. **(B)** Simplified diagram of dual vmThal-LEC/PER projection populations. Note that projections could be to either PER or LEC (magenta), but projections to PER are shown for simplicity.

While the present study identified dual vmThal-LEC/PER-projecting cell populations, there are several limitations to consider. First, this approach required stereotactic injections of CTB into two target sites with small enough volumes to be region specific, but large enough volumes to hit a significant proportion of the afferent synaptic terminal zones. Thus, while the presence of dual projecting cells is well concluded, this approach is not exhaustive of the combinations vmThal and LEC/PER subregions and there is a possibility that these populations could be further subdivided depending on where in vmThal and LEC/PER they project. Second, this study likely underestimates the actual proportion neurons that are dual projecting because the CTB infusions do not fully cover the target region, and because we used DAPI counts in our normalizations which include both glia and neurons. Future studies might better restrict the total cell count to only projection neurons since the long-range connectivity is of most interest in understanding the mPFC-HC communications. Likewise, better estimates also require more advanced cell counting and estimation procedures such as unbiased stereology. Lastly, it would be interesting to look at the calcium binding protein expression patterns in these populations, especially since these nicely divide up populations in vmThal and relate to their projection target profiles (Viena et al., 2021).

### 4.2. Potential roles for different dual projecting cell populations

These experiments identified three distinct cell populations from three different brain regions (MS, Sub, EC) with the potential to simultaneously influence the cortical and thalamic mPFC-HC communication routes. Anatomically, MS cells had the longest distance dual projections to both vmThal and LEC/PER, while vSub and EC cells were more local to LEC/PER in the sense that they are adjacent structures. Presumably each cell population contributes to the circuit differently.

The dual vmThal-LEC/PER-projecting MS cells are interesting because of their known role in hippocampal and medial entorhinal cortical theta activity, which is thought to synchronize and facilitate communication between mPFC and HC in service of learning, memory and decision making (Buzsáki, 2002; Dolleman-Van der Weel et al., 2019; Eichenbaum, 1996; Fuchs et al., 2016; Griffin, 2015; Schultheiss et al., 2020; Wirt & Hyman, 2017). For example, Zutshi et al., (2018) showed that HC theta power can be driven by long range MS cells, specifically GABAergic neurons. It is unknown here whether the dual projecting cells are GABAergic, glutamatergic and/or cholinergic cells, but this can be examined with immunohistochemical triple labeling approaches in future experiments. The dual vmThal-LEC/PER-projecting cells in MS, nonetheless, may drive theta synchrony in both mPFC-HC communication pathways helping to integrate multiple streams of information and orchestrating a large network of HC-cortical areas for advantageous memory-based decision making.

The dual vmThal-LEC/PER-projecting cell population probably contributes more as a major output structure of HC proper. The ventral subiculum in particular, is known to play a role in multiple learning and memory systems, as well as in spatial navigation (Canteras & Swanson, 1992; Ding et al., 2020; Ishihara et al., 2020; Ishizuka, 2001; Jay & Witter, 1991; Witter & Amaral, 2021). For instance, the subiculum may contribute to sharp wave-ripple (SWR) generation and the propagation (Imbrosci et al., 2021). It will be interesting to see if the dual vmThal-LEC/PER projecting cell population found in subiculum is related to SWR triggered/related activity in mPFC, and under what conditions this requires vmThal and LEC/PER pathways to be in sync. For example, the dual vmThal-LEC/PER-projecting cells found in vSub could be acting as a secondary SWR generator amplifying and expanding it to multiple systems, which may play an important role for the consolidation of detailed or contextualized information in memory.

The dual vmThal-LEC/PER-projecting cell population found in EC are in a unique position, a site of broad cortical convergence, to act as a representations communication hub between mPFC and HC (Agster et al., 2016; Beckstead, 1979; Burwell & Amaral, 1998b) facilitating or gating the directional flow of information. EC has been shown to be involved in multidimensional sensory representations and a primary input stream into and out of the hippocampus. Layer II/III specifically, was shown to be involved in temporal context memory (Suh et al., 2011), which requires mPFC-HC interactions (Eichenbaum, 2017; Jin & Maren, 2015; Sigurdsson & Duvarci, 2016; Vertes, 2006; Wirt & Hyman, 2017). Of note, most of the vmThal-projecting cells in EC were found in shallow layers II/III with few cells found in layers V, which is consistent with other reports (Dolleman-Van der Weel & Witter, 1996; Wouterlood, 1991; Wouterlood et al., 1990). Furthermore, the dual vmThal-LEC/PER-projecting cell population in EC was localized in layer II/III in both medial and lateral regions. This highly specific anatomy suggests the synchronizing outputs of EC, relating to both the cortical and thalamic pathways, are restricted to these layers.

### 4.3. Conclusion

In this current study, we presented evidence for three distinct cell populations within the mPFC-HC circuitry (MS, vSub, and EC) that project to both the thalamic (vmThal) and cortical (LEC/PER) communication hubs. Because the cortical and thalamic mPFC-HC connection routes have numerous contributions to learning, memory, and decision making (e.g., Anderson et al., 2016; Barker et al., 2007; Jayachandran et al., 2019), it is crucial to have an understanding how these different communication pathways are anatomically connected. Each population provides an opportunity for synchronous inputs to both the cortical and thalamic mPFC-HC communication routes, but each likely makes unique neurophysiological and behavioral contributions. However, future experiments are needed that are capable of isolating and manipulating each cell population individually, such as with a dual CRE-dependent retrograde AAV approach, to provide direct evidence of the contributions of these dual vmThal-LEC/PER-projecting cell populations.

## Abbreviations

ACC: Anterior cingulate cortex
CA1: Field of CA1 of Ammon’s horn
CA2: Field of CA2 of Ammon’s horn
CA3: Field of CA3 of Ammon’s horn
CTB: Cholera toxin subunit-B
DG: Dentate gyrus
dPL: Prelimbic cortex, dorsal
dSub: Subiculum, dorsal
IAM: Interanteromedial nucleus of thalamus
IL: Infralimbic cortex
LEC: Entorhinal cortex, lateral
LSd: Septum, lateral, dorsal
LSi: Septum, lateral, intermediate
LSv: Septum, lateral, ventral
MEC: Entorhinal cortex, medial
mPFC: Medial prefrontal cortex MS Septum, medial
PER: Perirhinal cortex
PT: Paratenial nucleus of thalamus
RE: Nucleus reuniens of thalamus
Rh: Rhomboid nucleus of thalamus
Sep: Septum nucleus
slm: Stratum lacunosum moleculare
sss: Superior sagittal sinus
Sub: Subiculum
vmThal: Ventral midline thalamus
vPL: Prelimbic cortex, ventral
vSub: Subiculum, ventral
3V: Third ventricle

## Acknowledgements

This work was supported by NIH grant R01 MH113626. A special thanks to all the members of the AllenLab, specifically our undergraduate research assistants, Sofia Leyva, and Victoria Blanco, who helped with all the cell counts, and proofreading. Also, we would like to thank the Animal Care Facility and Dr. Horatiu Vinerean DVM, DACLAM.

## Author Contributions

Conceptualization, T.A.A., M.J. and M.S.; Methodology, T.A.A., M.J. and M.S.; Investigation, M.S. and M.J.; Writing – Original Draft, M.S., M.J., G.E.R., and T.A.A.; Writing – Review and Editing, T.A.A., M.J., G.E.R. and M.S.; Funding Acquisition, T.A.A.; Resources, T.A.A.; Supervision, T.A.A.

